# Changes in levels of major yolk protein in the coelomic fluid and gonad during the reproductive cycle in wild sea urchins, *Mesocentrotus nudus*

**DOI:** 10.1101/241059

**Authors:** Kazuhiro Ura, Narumi Takei, Ichiro Higuchi, Tomoharu Yuhi, Osamu Nishimiya, Yasuaki Takagi

**Author notes:** Corresponding author: Kazuhiro Ura, Graduate School of Fisheries Sciences, Hokkaido University, 3-1-1 Minato-cho, Hakodate 041-8611, Japan.

## Abstract

Both female and male sea urchins accumulate the major yolk protein (MYP) in the nutritive phagocytes of immature gonads before gametogenesis, and MYP is the most abundant protein in the coelomic fluid of both sexes. In females, MYP in the coelomic fluid is taken up by the nutritive phagocytes and transported to the growing oocytes. This study examined quantitative changes of MYP in the coelomic fluid of both sexes during the reproductive cycle of wild sea urchins, *Mesocentrotus nudus*. Levels of MYP in the coelomic fluid of females increased and reached a peak at the histological pre-mature stage of gonad activity (i.e. Stage 3), and positive correlation between the MYP level and the gonadosomatic index (GSI) was observed. In male sea urchins the level of MYP in the coelomic fluid increased at the pre-mature stage, though positive correlation between the MYP level and GSI was not observed. These results indicate that MYP in the coelomic fluid is suitable as a biomarker of the onset and progression of sexual maturity in female sea urchins.

## Introduction

In sea urchins, the major yolk protein (MYP) is the most abundant yolk protein, and is stored in the yolk granules of eggs as a nutrient source during early development (Ozaki, 1980; Harrington and Easton, 1982; Kari and Rottmann, 1985), similar to the yolk proteins provided by vitellogenin in other species of oviparous animals (Sullivan et al., 2003; Hiramatsu et al., 2005). It has been reported that MYP is stored in nutritive phagocytes, which are versatile somatic cells, in the gonads of both sexes in sea urchins (Unuma et al., 1998). In males, the stored MYP is used as energy for spermatogenesis as gametogenesis proceeds. In females, some MYP is degraded and used as a nutrient source, while some is transported to the oocytes and actively packed into yolk granules through endocytosis via a dynamin-dependent mechanism (Unuma et al., 2003; Brooks and Wessel, 2004; Unuma et al., 2010).

MYP mRNA expression is ubiquitous in the digestive tract, coelomocytes and gonads of sea urchins (Unuma et al., 2001), while the major production sites of MYP are the digestive tract and gonads in adults of both sexes (Unuma et al., 2010). In both sexes of *Mesocentrotus nudus*, the MYP mRNA expression level increased during gonadal growth, and then decreased as gametogenesis proceeded in wild sea urchins (Ura et al., 2017). Whereas MYP is abundant in the coelomic fluid of male and female sea urchins, and is thought to be secreted in the digestive tract, as yet no histological analysis has provided evidence of this. Nonetheless, an *in vivo* experiment by Unuma et al. (2007) determined that MYP in the coelomic fluid is taken up by nutritive phagocytes in gonad, and this MYP is transported into the growing oocytes in females. To avoid confusion, we refer to MYP in coelomic fluid as CF-MYP, as originally reported by Unuma et al. (2007); we denote MYP synthesized in the gonads of both sexes as NP-MYP; finally, we use the term MYP in a broad sense when the type is not specified.

Fish vitellogenin is a female-specific protein and MYP precursor synthesized in the female liver, and vitellogenin is transported into the growing oocytes via the blood. Consequently, vitellogenin is used as a biomarker of the onset of puberty and the progression of sexual maturity in female fish (Hiramatsu et al., 2005; Mushirobira et al., 2013). However, we are not aware of any report using an assay system to profile CF-MYP during the reproductive cycle in adult sea urchins. Although CF-MYP is transported from coelomic fluid to the nutritive phagocytes of gonads and then into the growing oocytes, it is still unclear whether CF-MYP represents a suitable biomarker of the progression of sexual maturity in sea urchins. Thus, in the present study, we developed an assay system to examine changes in the levels of MYP in coelomic fluid and gonad, and changes in the level of total proteins in the coelomic fluid during the reproductive cycle in both sexes of wild sea urchins *(Mesocentrotus nudus)* collected at southern Hokkaido, Japan.

## MAterials and methods

### Animals and sampling

Sea urchins *Mesocentrotus nudus* were collected by diving at Usujiri in southern Hokkaido, Japan, from May 2015 to October 2016 (Ura et al., 2017). Sea urchins were transported to the Faculty of Fisheries Sciences of Hokkaido University, where the gonads were excised and weighed. The gonadosomatic index (GSI) was calculated for each animal as follows: GSI (%) = 100 × wet weight of gonad/total wet body weight. A small portion of each gonad sampled was fixed in Bouin’s solution for histological examination of the reproductive stage, and the reminder was stored at −30°C until analysis.

After removing the peristomial membrane, coelomic fluid was collected and combined with an equal volume of anticoagulant solution (20 mmol l^−1^ Tris-HCl, 500 mmol l^−1^ NaCl, 30 mmol l^−1^ EDTA; pH 7.4) and then centrifuged at 900 *g* for 10 min at 4°C to separate the coelomic fluid, according to the method of Schillaci et al. (2013). The coelomic fluids were stored at −30°C until analysis.

### Histology of gonads

The tissue samples were dehydrated through a graded ethanol series and embedded in paraffin; 6-μm-thick serial sections were mounted on glass slides and stained with hematoxylin and eosin. The gonadal maturity of each animal was classified according to the five stages described by Unuma et al. (1996): Stage 1 (recovering), Stage 2 (growth), Stage 3 (pre-mature), Stage 4 (mature), and Stage 5 (spent). The gonads of 15–30 individuals were collected and analyzed each month.

### Purification of NP-MYP

The immature male gonads were homogenized with five volumes of 10 mmol l^−1^ Tris-HCl (pH 8.0), containing 10 mmol l^−1^ NaCl, 1 mmol l^−1^ EDTA, and 0.1% NaN_3_, using a Teflon homogenizer. The homogenate was centrifuged at 17,000 *g* for 20 min at 4°C, and supernatant was collected. Ion-exchange chromatography was performed on the gonad extract using a 2.5 × 8-cm column of DEAE cellulose (TOYOPEAL DEAE) equilibrated with 50 mmol l^−1^ Tris-HCl buffer (pH 7.5). The retained proteins were eluted in stepwise fashion using the same buffer at various molar concentrations of NaCl (0, 100, 200 and 300 mmol l^−1^) at 4°C. Fractions (100 mmol l^−1^ NaCl) rich in the targeted protein were pooled and dialyzed against 20 mmol l^−1^ KP at 4°C for overnight. The dialyzed sample was fractionated by hydroxylapatite chromatography, and MYP was eluted at 400 mmol l^−1^ KP. This fraction was concentrated by ultrafiltration to 2 ml and fractionated twice by Superose 6 10/300 GL gel filtration column (GE Healthcare Life Sciences, Little Chalfont, UK) equilibrated with 50 mmol l^−1^ Tris-HCl buffer (pH 7.5) containing 500 mmol l^−1^ NaCl. The single protein peak was the target MYP. At each purification step, electrophoresis and immunological procedures were used to ensure that the target protein was present. Purified MYP concentration was measured as described by Lowry et al. (1951), with bovine serum albumin (BSA) (Bio-Rad Laboratories Inc., Hercules, CA, USA) as the standard.

### Preparation of antisera

A polyvalent antiserum against gonad extract proteins (anti-gonad) was raised in rabbits. The specific antiserum to purified NP-MYP (anti-MYP) was obtained from a rabbit immunized with 1 ml of solution containing 300 μg of purified NP-MYP mixed with an equal volume of Freund’s complete adjuvant (Wako Pure Chemical Industries, Osaka, Japan). The rabbit received four such immunizations at about 7-day intervals. After four injections, blood was collected from the ear vein of the rabbit. The blood was allowed to clot for about 1 h at room temperature, and was then stored overnight at 4°C. The blood was centrifuged at 17,000 *g* for 15 min at 4°C, and the supernatant was collected as antiserum.

### Electrophoresis and immunological procedure

Disc-polyacrylamide gel electrophoresis (disc-PAGE) was carried out in 5% polyacrylamide gel using a Tris-glycine buffer system, following Davis (1964). The gel was stained with 1% Amido black 10B in 7% acetic acid and destained with 7% acetic acid. Immunoelectrophoretic analysis was performed with 1% agarose in 190 mmol l^−1^ Tris-HCl buffer (pH 8.6). The immunoprecipitates were stained with 1% Amido black 10B in 7% acetic acid and destained with 7% acetic acid in the dried gel. Double immunodiffusion using anti-MYP was performed in 1% agarose gel, using the method of Ouchterlony (1953).

### Assay of MYP

Single radial immunodiffusion (SRID) was carried out according to the procedure of Mancini et al. (1965). Antiserum to purified MYP was diluted at 56°C in a solution of 1% (w/v) agarose (Nacalai, HGT) in 190 mmol l^−1^ Tris-HCl buffer (pH 8.6). Next, 15 ml of the hot solution was layered onto a 10 × 10 cm sheet of GelBond film (GE Healthcare Life Sciences). The SRID plate was incubated in a moist chamber at room temperature for 2 days; after incubation, it was washed with 0.9% NaCl, dried on filter paper, stained with 1% Amido black 10B in 7% acetic acid, and destained with 7% acetic acid. Purified MYP (25, 50, 100, 200 and 400 μg ml^−1^) was used as the standard for quantitative SRID in the gonads. Purified MYP (20, 30, 60, 120 and 200 μg ml^−1^) was used as the standard for quantitative SRID in the coelomic fluids.

### Measurement of total protein concentration in the coelomic fluid

The total protein concentrations in the coelomic fluids were measured as described by Lowry et al. (1951), with BSA as the standard.

### Statistical analysis

Data were expressed as mean ± SE. Statistical differences between means across the stages of gonad activity were determined by one-way ANOVA and subsequent Dunnett’s test, where Stage 1 (recovering) was used as a control group. Statistical significance is denoted at the level *P* < 0.01 (*) or *P* < 0.05 (**). Correlation was determined by Spearman’s rank correlation test.

## Results

### Changes in GSI during the reproductive cycle

The change in the GSI during the reproductive cycle of *Mesocentrotus nudus* is shown in Fig. 1. In the wild population of sea urchins, the mean value significantly increased from 9.42 ± 0.37% at Stage 1 (recovering) to 13.02 ± 0.59 at Stage 2 (growth) and 12.18 ± 0.56 at Stage 3 (pre-mature), and thereafter decreased at Stage 4 (mature) (Fig. 1A). In females, the mean value significantly increased from 9.67 ± 0.53% at Stage 1 to 14.28 ± 0.74% at Stage 2, and thereafter gradually decreased (Fig. 1B). In males, the mean value significantly increased from 9.06 ± 0.48% at Stage 1 to 12.02 ± 0.66% at Stage 3, and then decreased to 7.35 ± 1.22% at Stage 4 (Fig. 1C).

**Figure 1.**
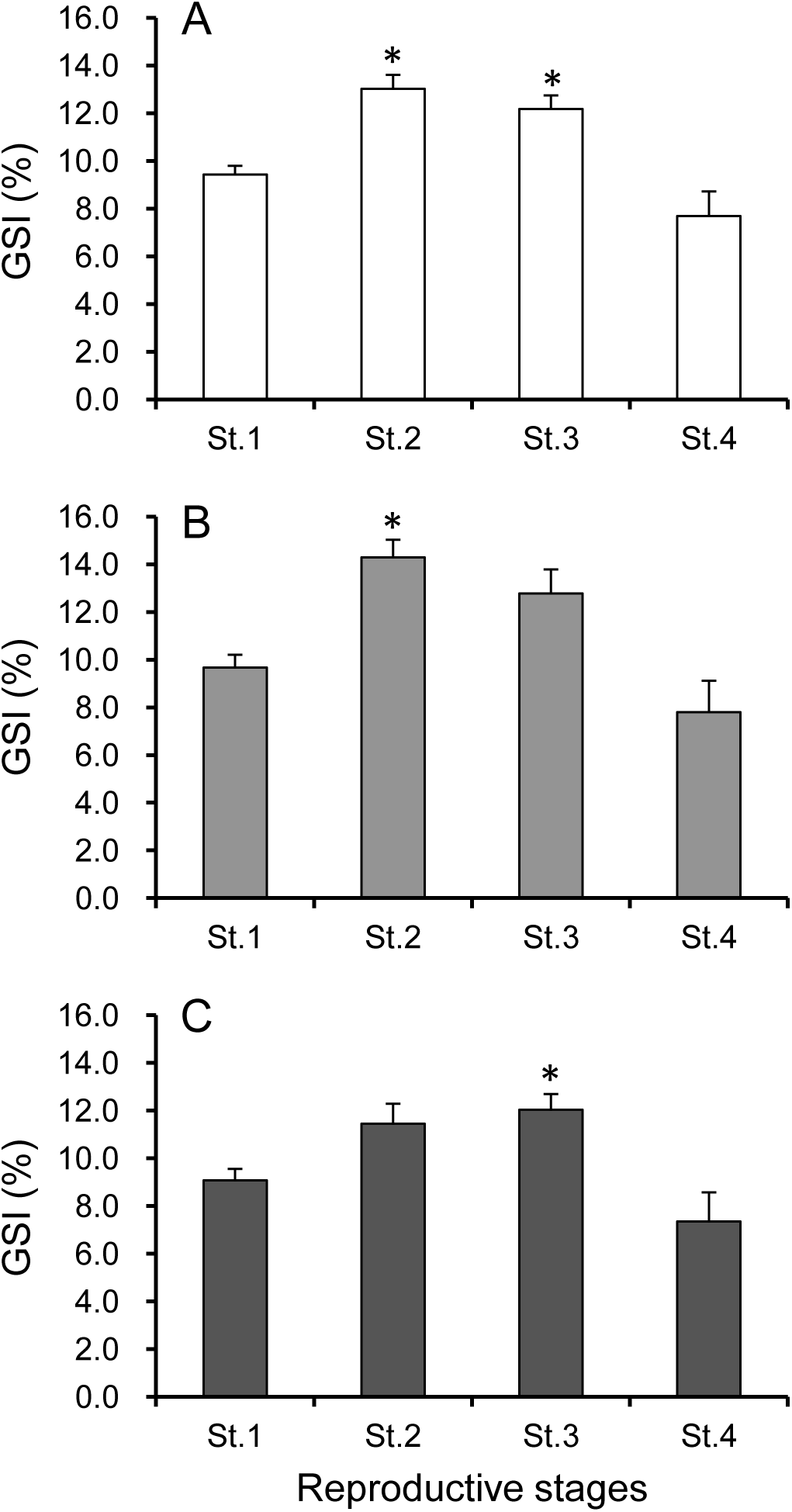
Changes in gonadosomatic index (GSI) in the wild population (A) and in female (B) and male (C) sea urchins at different reproductive stages. Values are mean ± SE. Significant differences between means at the level *P* < 0.01 (*) were determined using Dunnett’s test, where Stage 1 (St. 1) gonad activity (i.e. recovering) was used as a control group.

### Purity of the isolated NP-MYP

The purity of the isolated NP-MYP preparation was assessed by means of disc-PAGE and immunoelectrophoresis using a polyvalent antiserum to gonad extract and a specific antiserum to NP-MYP. One homogenous band was observed on disc-PAGE when stained with Amido black 10B (Fig. 2A). In immunoelectrophoresis, the purified NP-MYP produced a single precipitin line against the polyvalent antiserum to gonad extract proteins. Conversely, the antiserum raised against the purified NP-MYP developed only a single precipitin line with male gonad extract protein (Fig. 2B).

**Figure 2.**
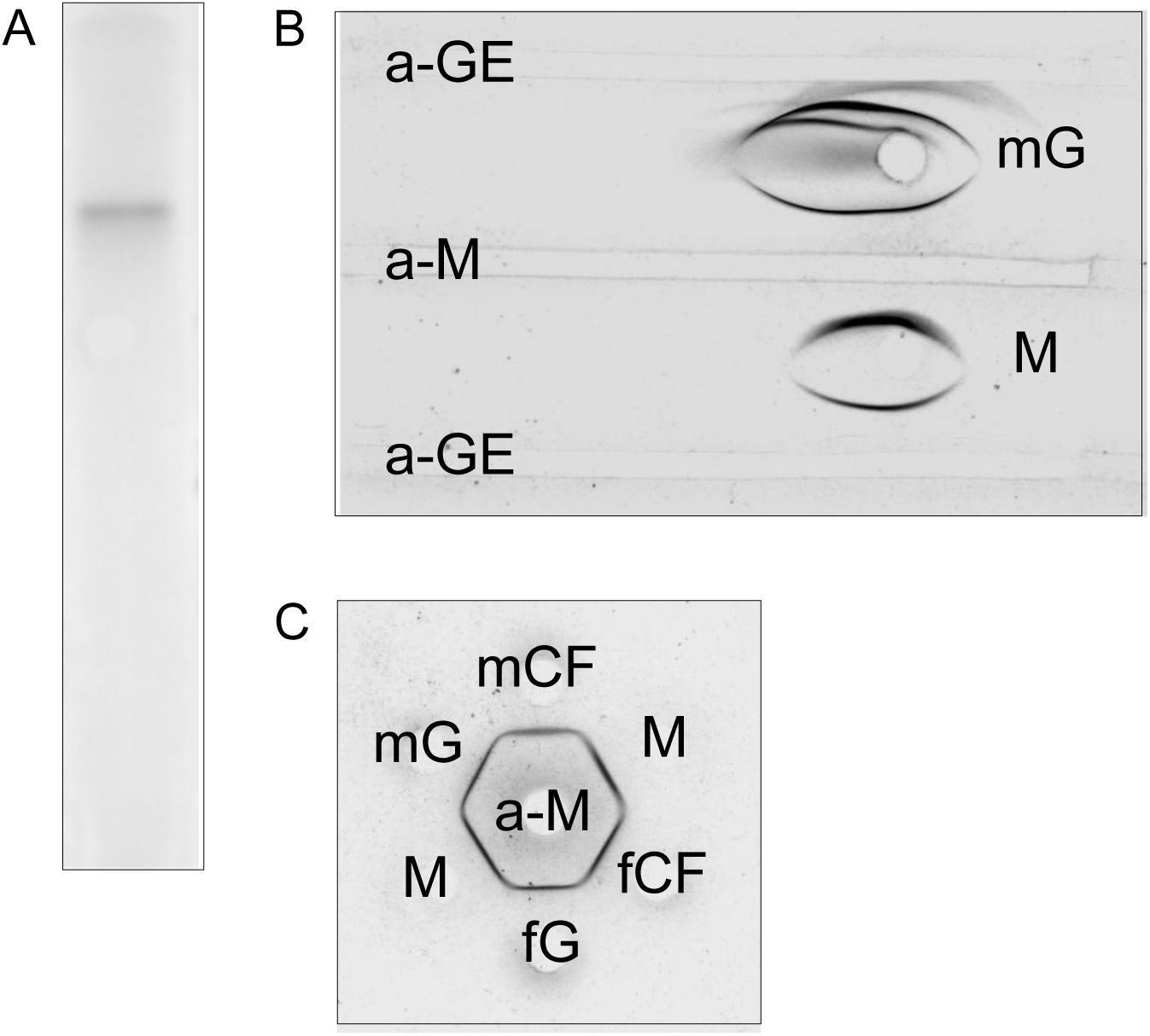
Observations of disc electrophoresis (A) and immunoelectrophoresis (B) of purified NP-MYP from immature male gonads, and the precipitin reaction of MYP in double immunodiffusion (C). Disc electrophoresis was stained with Amido black 10B; a-GE: polyvalent antiserum to gonad extracts; a-M: specific antiserum to purified NP-MYP; M: purified NP-MYP; mG: male gonad extract; fG: female gonad extract; mCF: male coelomic fluid; fCF: female coelomic fluid.

### Antigenic relationships between MYP in coelomic fluid and gonad

The result of double immunodiffusion of coelomic fluids and gonad extract protein using antiserum against NP-MYP is shown in Fig. 2C. A precipitin line of coelomic fluids of both sexes appeared to generate a fuse against purified NP-MYP from male gonads; and a precipitin line of gonad extract protein of both sexes appeared to generate a fuse against purified NP-MYP.

### Quantitative measurement of MYP in gonads

Different dilutions of the antiserum to MYP were incorporated into the agarose gel used for SRID. A concentration of 2% antiserum gave the best quantitative results for measurement of MYP in the gonads of both sexes (Fig. 3A). Using this dilution, the squared diameter of a precipitate ring was directly proportional to the amount of the sample, in the range of 25, 50, 100, 200 and 400 μg ml^−1^ standards, as shown in Fig. 3B. The diameter of precipitation rings produced by purified NP-MYP were directly proportional to the concentration of the MYP standard (R^2^ = 0.9927), and the serial dilutions of gonad samples ran parallel to the standard curve. The inter-assay coefficient of variation was 2.55% (n = 20) and the intra-assay coefficient of variation was 0.87% (n = 4). Recovery of various concentrations (25, 50, 100, 200 and 400 μg ml^−1^) of purified NP-MYP standard added to gonad extract was from 93.8 to 98.3%.

**Figure 3.**
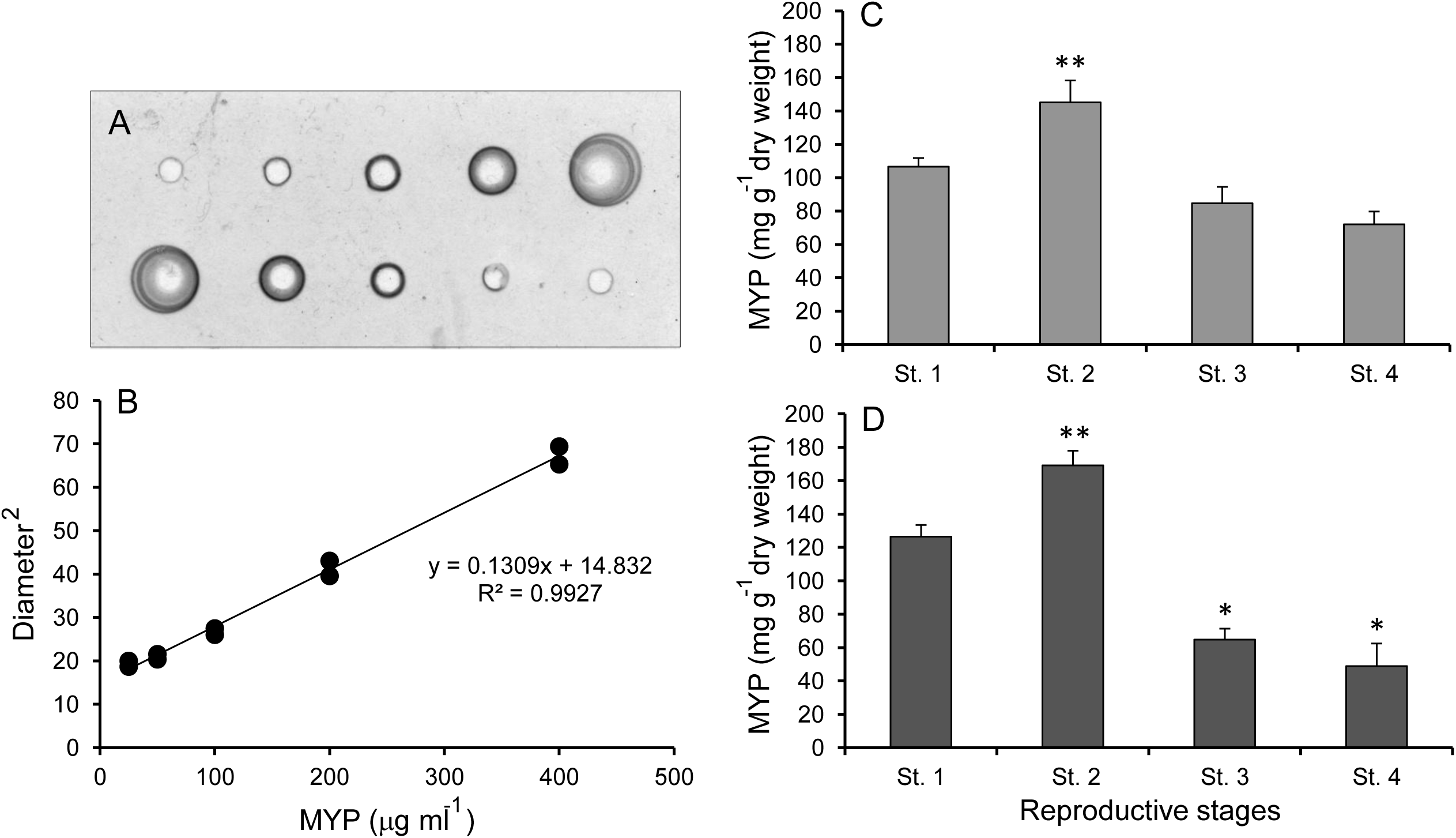
A single radial immunodiffusion (SRID) plate with 2% concentration of antiserum to NP-MYP (A) and their standard curve (B) for measurement of MYP in the gonads of both sexes of sea urchin. Quantitative changes in the content of MYP in the gonad of females (C) and males (D) at different reproductive stages. Values are mean ± SE. Significant differences between means at the level *P* < 0.01 (*) or *P* < 0.05 (**) were determined using Dunnett’s test, where Stage 1 (St. 1) gonad activity (i.e. recovering) was used as a control group.

The concentration of MYP in the gonads of both sexes of *M. nudus* was measured during the reproductive cycle (Fig. 3C, D). In females, the mean levels of MYP during Stages 1, 2, 3 and 4, respectively, were: 106.5 ± 5.2 mg g^−1^ (dry weight) (n = 6), 145.1 ± 13.1 mg g^−1^ (dry weight) (n = 6), 84.6 ± 9.9 mg g^−1^ (dry weight) (n = 6), and 72.0 ± 7.7 mg g^−1^ (dry weight) (n = 6) (Fig. 3C). In males, the mean levels of MYP were: 126.4 ± 7.0 mg g^−1^ (dry weight) (n = 5), 169.0 ± 8.8 mg g^−1^ (dry weight) (n = 6), 64.8 ± 6.5 mg g^−1^ (dry weight) (n = 6), and 48.8 ± 13.6 mg g^−1^ (dry weight) (n = 3) (Fig. 3D). Moreover, significant changes during the reproductive cycle were found for both females and males. In females, the level of MYP significantly increased from Stage 1 to Stage 2 (*P* < 0.05) and then gradually decreased at Stage 4. In males, the level of MYP significantly likewise increased from Stage 1 to Stage 2 (*P* < 0.05), but then drastically decreased during the rest of the reproductive cycle.

### Quantitative measurement of MYP in coelomic fluids

A concentration of 0.5% antiserum gave the best quantitative results for measurement of MYP in the coelomic fluids of both sexes (Fig. 4A). Using this dilution, the squared diameter of a precipitate ring was directly proportional to the amount of the sample, in the range of the 20, 30, 60, 120 and 200 μg ml^−1^ standards, as shown in Fig. 4B. The diameter of the precipitation rings produced by purified NP-MYP were directly proportional to the concentration of the MYP standard (R^2^ = 0.993), and the serial dilutions of coelomic-fluid samples ran parallel to the standard curve. The inter-assay coefficient of variation was 3.81% (n = 20) and the intra-assay coefficient of variation was 1.79% (n = 4). Recovery of various concentrations (20, 30, 60, 120 and 200 μg ml^−1^) of purified NP-MYP standard added to coelomic fluid was from 96.9 to 99.9%.

**Figure 4.**
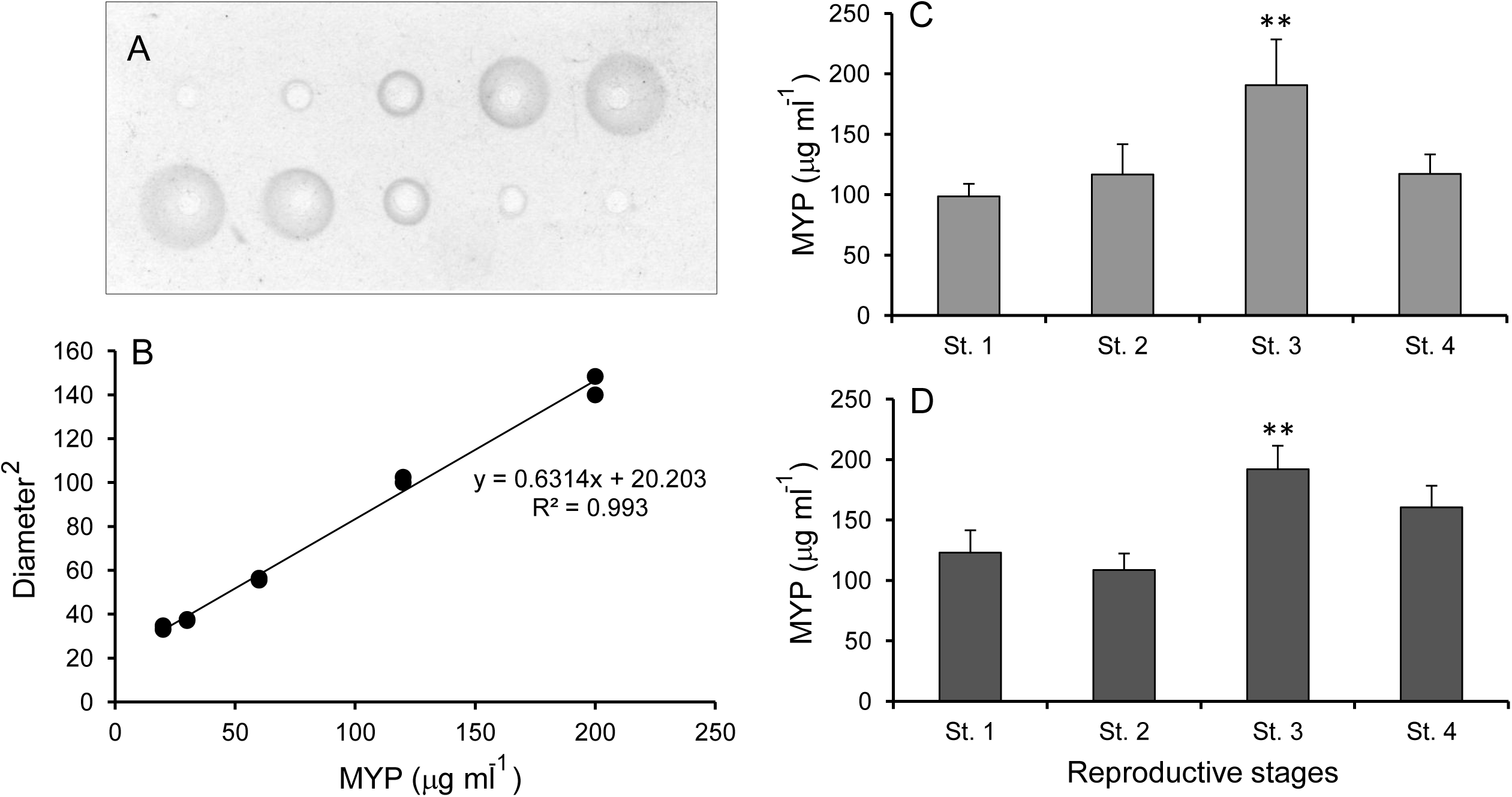
SRID plate with 0.5% concentration of antiserum to NP-MYP (A) and theirStandard curve (B) for measurement of MYP in the coelomic fluids of both sexes of sea urchin. Quantitative changes in the content of CF-MYP in the gonads of females (C) and males (D) at different reproductive stages. Values are mean ± SE. Significant differences between means at *P* < 0.05 (**) were determined using Dunnett’s test, where Stage 1 (St. 1) gonad activity (i.e. recovering) was used as a control group.

The concentration of MYP in the coelomic fluids of both sexes of *M. nudus* was measured during reproductive cycle (Fig. 4C, D). In females, the mean levels of MYP for Stages 1, 2, 3 and 4, respectively, were: 98.6 ± 10.3 μg ml^−1^ (n = 10), 116.6 ± 25.0 μg ml^−1^ (n = 10), 190.6 ± 37.8 μg ml^−1^ (n = 7), and 117.1 ± 16.1 μg ml^−1^ (n = 10) (Fig. 4C). In males, the mean levels of MYP for Stages 1, 2, 3 and 4, respectively, were: 122.9 ± 18.6 μg ml^−1^ (n = 10), 108.6 ± 13.6 μg ml^−1^ (n = 10), 192.0 ± 19.4 μg ml^−1^ (n = 10), and 160.5 ± 17.8 μg ml^−1^ (n = 3) (Fig. 4D). In both females and males, the level of CF-MYP significantly increased from Stage 1 to Stage 3 (*P* < 0.05) and then gradually decreased at Stage 4.

### Correlation between GSI and CF-MYP level

Figure 5 shows the correlation between GSI values and CF-MYP levels in female and male sea urchins. In females (n = 37), the correlation coefficient was R^2^ = 0.2483, and a significant positive correlation was observed (*P* = 0.0018). In males (n =33), the correlation coefficient was R^2^ = 0.0019, but no significant positive correlation was observed (*P* = 0.6519).

**Figure 5.**
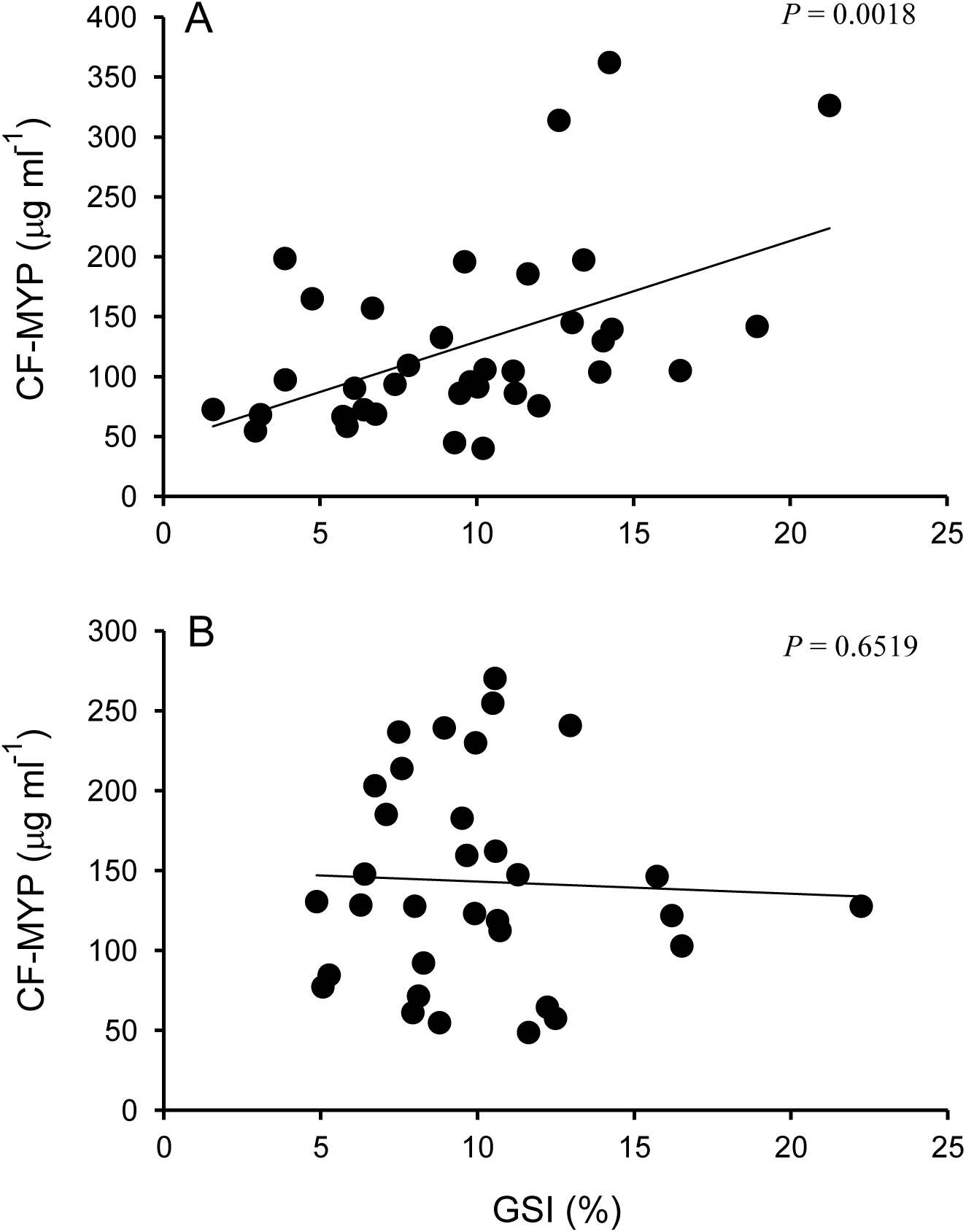
Correlation of gonadosomatic index (GSI) with MYP in the coelomic fluid of female (A) and male (B) sea urchins.

### Changes in levels of total proteins and the ratio of MYP in coelomic fluids

The concentration of total proteins in the coelomic fluids was measured during the reproductive cycle (Fig. 6). In the wild population, the mean value significantly increased from 0.337 ± 0.015 mg ml^−1^ (n = 20) at Stage 1 to 0.427 ± 0.025 mg ml^−1^ (n = 17) at Stage 3, and then decreased to 0.362 ± 0.021 mg ml^−1^ (n = 13) at Stage 4 (Fig. 6A). In females, the mean levels of total protein were 0.328 ± 0.023 mg ml^−1^ (n = 10), 0.373 ± 0.030 mg ml^−1^ (n = 10), 0.431 ± 0.038 mg ml^−1^ (n = 7) and 0.342 ± 0.022 mg ml^−1^ (n = 10) for Stages 1, 2, 3 and 4, respectively (Fig. 6B). In males, the mean levels of total protein were 0.345 ± 0.021 mg ml^−1^ (n = 10), 0.354 ± 0.034 mg ml^−1^ (n = 10), 0.424 ± 0.033 mg ml^−1^ (n = 10) and 0.427 ± 0.037 mg ml^−1^ (n = 3) for Stages 1, 2, 3 and 4, respectively (Fig. 6C). Although significant changes were not observed during the reproductive cycle in either the females or males, the mean levels increased slightly from Stage 1 to Stage 3 in both sexes.

**Figure 6.**
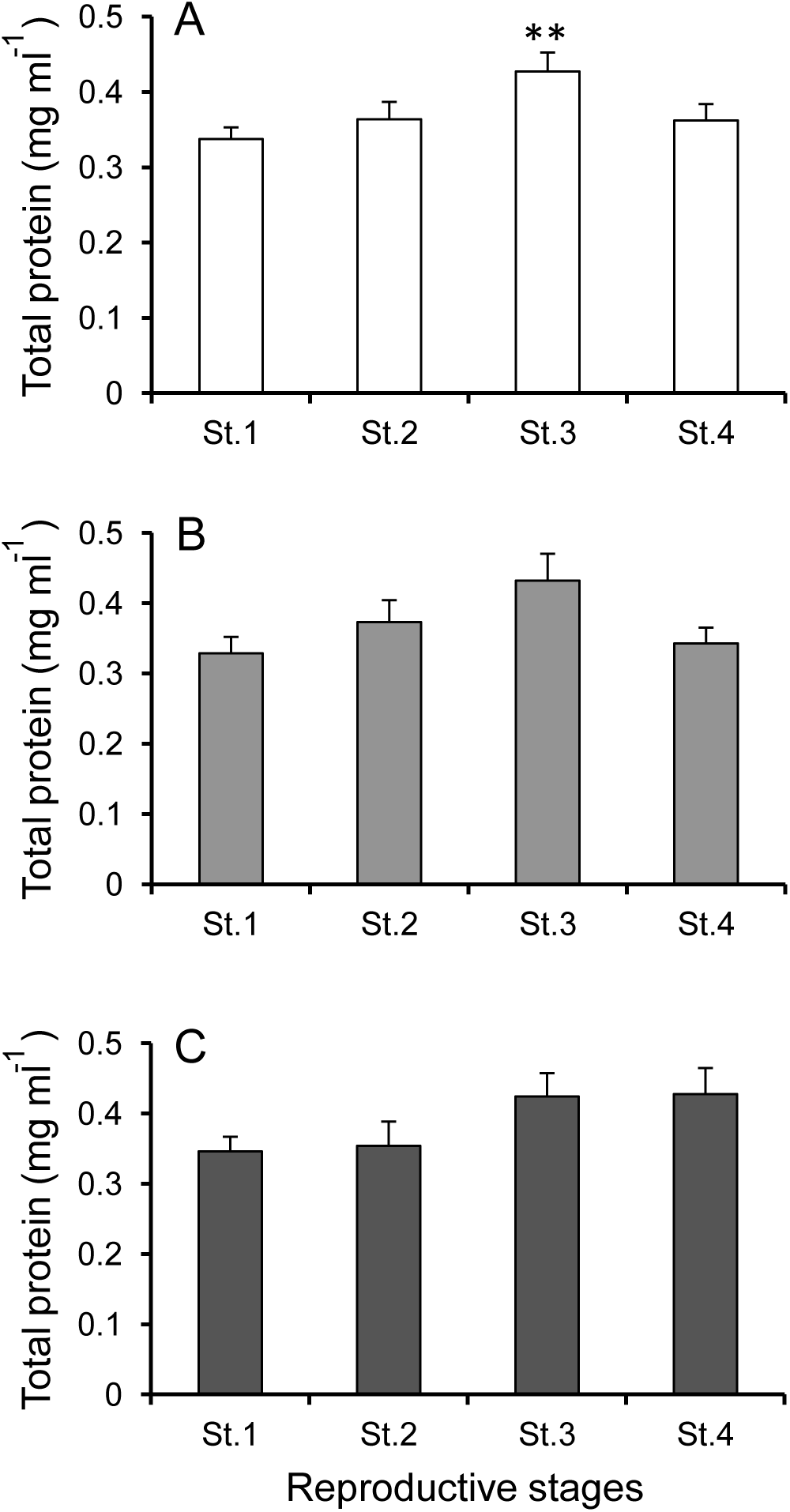
Quantitative changes in the content of total proteins in the coelomic fluid in the wild population (A), and in females (B) and males (C) at different reproductive stages. Values are mean ± SE. Significant differences between means at *P* < 0.05 (**) were determined using Dunnett’s test, where Stage 1 (St. 1) gonad activity (i.e. recovering) was used as a control group.

### Correlation between GSI and total protein level in coelomic fluids

Figure 7 shows the correlation between GSI and total protein level in the coelomic fluids. In the wild population (n = 70), the correlation coefficient was R^2^ = 0.086, and a significant positive correlation was observed (*P* = 0.031) (Fig. 7A). The correlation coefficient was R^2^ = 0.382, and a significant positive correlation was observed (*P* = 0.00014) in females (n = 37); however, a positive correlation was not observed (*P* = 0.425) between GSI and total protein level in males (n = 33) (Fig. 7B, C).

**Figure 7.**
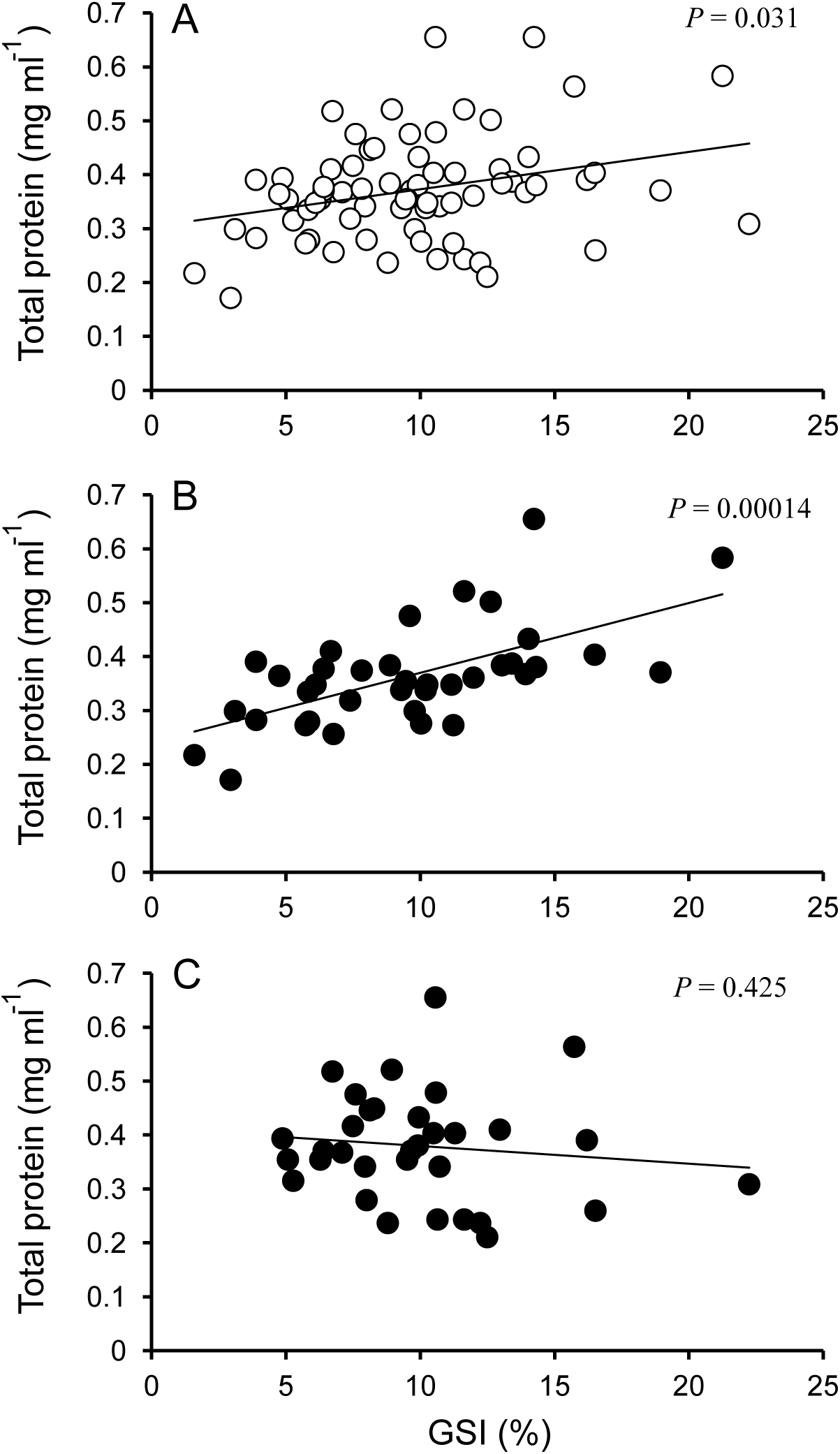
Correlation of gonadosomatic index (GSI) with total protein levels in the coelomic fluid of the wild population (A) and in female (B) and male (C) sea urchins.

## Discussion

This study aimed to identify whether CF-MYP is useful as a biomarker of the progression of sexual maturity in sea urchins. Thus, we examined changes in the concentration of MYP in the gonads and coelomic fluids of male and female *Mesocentrotus nudus* over the course of the reproductive cycle. We used wild sea urchins collected from southern Hokkaido, Japan, since we previously conducted a detailed study of the species’ reproductive cycle and examined transcription-level changes of the MYP in gonads of both sexes during the different reproductive stages (Ura et al., 2017).

The GSI significantly increased at Stage 2 (growth) and Stage 3 (pre-mature) in the wild population (Fig. 1A). The GSI increased from Stage 1 (recovering) to Stage 2 or Stage 3 in females and males, and then that gradually decreased in Stage 4 (mature) (Fig. 1B, C). In wild sea urchins *Paracentrotus lividus* collected off southern France the GSI peaked at Stage 3 and thereafter decreased (Spirlet et al., 1998). Agatsuma et al. (1988) likewise examined wild *M. nudus* (previously known as *Strongylocentrotus nudus)* from southern Hokkaido, and similarly reported that GSI peaked around Stage 3, and then decreased during the spawning season.

To develop an assay system to measure MYP in sea urchin, we performed purification of NP-MYP and obtained a specific antiserum against NP-MYP. Purity of the NP-MYP was assessed by disc-PAGE and immunoelectrophoresis. The purified NP-MYP yielded one band in disc-PAGE and a single precipitin line when reacted against a polyvalent antiserum to gonad extract as well as when it was precipitated by specific antiserum to NP-MYP. These results indicate that the NP-MYP preparation was electrophoretically and immunologically pure (Fig. 2A, B). The MYP stored in the nutritive phagocytes of sea urchin gonads is understood to be a glycoprotein (Ozaki et al., 1986). Furthermore, the antigenic relationship between gonad extracts and coelomic fluids of both sexes in double immunodiffusion indicated that the antigenicity of MYP in gonads was immunologically the same as MYP in the coelomic fluids of both sexes (Fig. 2C). These results demonstrate that this specific antiserum against NP-MYP can be used to measure MYP levels in the gonads and coelomic fluids of female and male sea urchins.

In the present study, changes in the concentration of MYP in the gonads of both sexes during the reproductive cycle were determined by SRID using the specific antiserum raised against the purified NP-MYP (Fig. 3). In female gonad, concentrations of MYP peaked at Stage 2 and gradually decreased as gametogenesis proceeded. In male gonad, concentrations of MYP likewise peaked at Stage 2 but decreased to rapidly Stage 4. This profile of MYP content in the gonads of both sexes is similar to that previously reported for cultured *Pseudocentrotus depressus* (Unuma et al. 2003). In our previous study of *M. nudus* (Ura et al., 2017), MYP mRNA expression levels significantly increased from Stage 1 and reached a peak at Stage 2; thereafter the levels gradually decreased in females, but drastically decreased in males. These results suggest that MYP in the gonads of both sexes of sea urchin is synthesized and stored at Stage 2; thereafter, a part of the MYP stored in the nutritive phagocytes is transported into growing oocytes in females, while MYP functions as a nutrient for gametogenesis in males.

MYP is an abundant protein in the coelomic fluid of both sexes in sea urchins (Giga and Ikai, 1985; Unuma et al., 1998), and MYP mRNA is expressed in the digestive tract, gonads and coelomocytes (Unuma et al., 2001). The coelomocytes of sea urchins are classified as four types: phagocytes, vibratile cells, and red and white morula cells (Bertheussen and Seljelid, 1978; Gerardi et al., 1990; Unuma et al., 2010). Unuma et al. (2010) observed the expression of MYP mRNA mainly in vibratile cells and white morula cells in sea urchin. Without anticoagulant solution, vibratile cells become immediately broken *in vitro* (Matsutani, 1995). This suggested the importance of collecting the coelomic fluid with an anticoagulant solution to determine CF-MYP concentrations in sea urchins, as carried out in the present study. Accordingly, we examined changes in the concentration of CF-MYP during the reproductive cycle in both males and females. Furthermore, we were aware of no previous reports using an assay system for profiling CF-MYP during the reproductive cycle in adult sea urchins. In this study, changes in the concentration of MYP in the coelomic fluids of both sexes during the reproductive cycle were determined by SRID (Fig. 4), and significant changes were found for both female and male sea urchins. A positive correlation between CF-MYP levels and GSI was observed in females, but not in males (Fig. 5). In an exquisitely designed *in vivo* experiment, Unuma et al. (2007) found that CF-MYP is taken up by nutritive phagocytes in the gonads, and is finally transported into the growing oocytes in females. In teleost fishes, the level of serum vitellogenin (an MYP precursor) showed a significant positive correlation with GSI (Mushirobira et al., 2013). These results indicate that CF-MYP is suitable as a biomarker of the onset of puberty and the progression of sexual maturity in female sea urchins, similar to vitellogenin in fishes. Whereas, in males, the level of CF-MYP significantly increased and reached a peak at Stage 3, it did not positively correlate with GSI; therefore, CF-MYP is not suitable as a biomarker of the progression of sexual maturity in male sea urchins. It is generally understood that MYP stored in the male gonad decreases with maturation, a phenomena observed in the present study. However, further investigation is needed to explain the high levels present in males at Stage 3, and to exactly determine the role of CF-MYP in male sea urchins.

We also examined changes in the concentration of total proteins in the coelomic fluids of both sexes of wild sea urchin during the reproductive cycle. In the wild population, total protein concentration increased from Stage 1 and peaked at Stage 3, and thereafter decreased to a low level at Stage 4 (Fig. 6). In a natural population of *Strongylocentrotus purpuratus* the concentration of total proteins in the coelomic fluid increased with an increasing GSI, and then decreased at a mature stage, with total protein concentration in the coelomic fluid reported as 0.20–0.45 mg ml^−1^ (Holland et al., 1967). In this study, the concentration of total proteins in the coelomic fluids were 0.17–0.65 mg ml^−1^ in wild *M. nudus*, a range similar to that found for purple sea urchin as reported by Holland et al. (1967). However, the observed changes in the concentration of total proteins in the coelomic fluids of females and males were not significant. A significant positive correlation between total protein level and GSI was observed for females, but not for males. Moreover, the strong positive correlation between total protein level and GSI in females, and not between CF-MYP and GSI, suggests that CF-MYP and another protein increases as gametogenesis proceeds in females. However, the identification and characterization of proteins in the coelomic fluids of sea urchins before maturity of the gametes remains to be done.

In conclusion, CF-MYP appears suitable as a biomarker of the onset of puberty and the progression of sexual maturity in female sea urchins, but not in males. This finding indicates that CF-MYP is a part of the MYP stored in the eggs of sea urchins.

## Acknowledgments

We thank Dr Hiroyuki Munehara (Usujiri Fisheries Station, Field Science Center for the Northern Biosphere, Hokkaido University, Japan) for providing samples of sea urchin gonads and for helpful advice. Cynthia Kulongowski (MSc), with the Edanz Group (www.edanzediting.com/ac), edited a draft of this manuscript. The present study was supported by a Grant-in-Aid for Scientific Research from the Ministry of Education, Culture, Sports, Science and Technology, Japan (No. 25660159), and the Science and Technology Research Promotion Program for Agriculture, Forestry, Fisheries and Food Industry.

